# Investigating genomic prediction strategies for grain carotenoid traits in a tropical/subtropical maize panel

**DOI:** 10.1101/2023.12.29.573624

**Authors:** Mary-Francis LaPorte, Willy B. Suwarno, Pattama Hannok, Akiyoshi Koide, Peter Bradbury, José Crossa, Natalia Palacios-Rojas, Christine Helen Diepenbrock

## Abstract

Vitamin A deficiency remains prevalent on a global scale, including in regions where maize constitutes a high percentage of human diets. One solution for alleviating this deficiency has been to increase grain concentrations of provitamin A carotenoids in maize (*Zea mays* ssp. *mays* L.)—an example of biofortification. The International Maize and Wheat Improvement Center (CIMMYT) developed a Carotenoid Association Mapping panel of 380 inbred lines adapted to tropical and subtropical environments that have varying grain concentrations of provitamin A and other health-beneficial carotenoids. This project assesses the accuracy of several genomic prediction (GP) strategies for these traits (β-carotene, β-cryptoxanthin, provitamin A, lutein, and zeaxanthin) within and between four environments in Mexico. Methods employing Ridge Regression-Best Linear Unbiased Prediction, Elastic Net, or Reproducing Kernel Hilbert Spaces had high accuracy in predicting all tested provitamin A carotenoid traits and outperformed Least Absolute Shrinkage and Selection Operator. Furthermore, prediction accuracies were higher when using genome-wide markers rather than only the markers proximal to two previously identified carotenoid-related genes that have been used in marker-assisted selection, suggesting that GP is worthwhile for these traits, even though key genes have already been identified. Prediction accuracy was maintained for all traits (except lutein) in between-environment prediction. The TASSEL (Trait Analysis by aSSociation, Evolution, and Linkage) Genomic Selection plugin performed as well as other more computationally intensive methods for within-environment prediction. The findings observed herein indicate the utility of GP methods for these traits and could inform their resource-efficient implementation in biofortification breeding programs.

## Introduction

Vitamin A deficiency is a global issue, impacting approximately 30% of children less than five years old worldwide, that can cause xerophthalmia (conjunctival and corneal dryness), blindness (particularly night blindness), morbidity, and mortality, particularly for children and pregnant women (World Health Organization 2009, 2014; Wirth et al. 2017; Hodge and Taylor 2023). Deficiencies can be alleviated through increased intake of provitamin A carotenoids in the diet (Bouis and Welch 2010; Pixley et al. 2013; Prasanna et al. 2020). Biofortification is a multifaceted strategy to address malnutrition through the enhancement of nutritional quality in crops via breeding and/or agronomic practices (Bouis and Welch 2010). Biofortifying maize with higher grain concentrations of provitamin A carotenoids through breeding can help alleviate Vitamin A deficiency, especially in regions, including parts of sub-Saharan Africa, where a large portion of the daily diet typically comes from maize (*Zea mays*) (Bouis and Welch 2010; Pixley et al. 2013; Saltzman et al. 2013).

Ongoing biofortification efforts have resulted in lines with higher concentrations of provitamin A carotenoids (12-15 μg/g dry weight) than the average for maize grain (<1.5 μg/g dry weight) (HarvestPlus, International Center for Tropical Agriculture (CIAT), Cali, Colombia and Andersson 2017; Ortiz et al. 2018). The International Maize and Wheat Improvement Center (CIMMYT, a member of CGIAR), developed and used a Carotenoid Association Mapping (CAM) panel of 380 maize inbred lines with different levels of grain carotenoids with growth phenotypes adapted to tropical and subtropical environments, which are relevant to the global target regions for carotenoid biofortification in sub-Saharan Africa and Latin America (Suwarno et al. 2015). The grain carotenoid traits of interest for the present study—β-carotene, β-cryptoxanthin, lutein, zeaxanthin, and provitamin A—were found to be highly heritable in the CAM panel in three environments, with broad-sense heritabilities ranging from 0.89 to 0.93 as reported by Suwarno et al. (2015). Ten lines included in the CAM panel have been biofortified (using marker-assisted selection; MAS) for an allele of *crtRB1 (β-carotene hydroxylase 1)* previously found to be associated with higher levels of provitamin A carotenoids (Yan et al. 2010; Babu et al. 2013; Suwarno et al. 2015).

Provitamin A and other health-beneficial carotenoids are synthesized in maize grain, primarily in the endosperm (Blessin et al. 1963; Weber 1987). Carotenoids are C40 isoprenoids, which can be classified as carotenes (including β-carotene) and xanthophylls (oxygenated carotenoids, including β-cryptoxanthin, zeaxanthin, and lutein). β-carotene and β-cryptoxanthin are the provitamin A carotenoids included in this study. Provitamin A is a derived trait, calculated based on retinol equivalents (Babu et al. 2013, Suwarno et al. 2015); i.e., provitamin A = β-carotene + 0.5(β-cryptoxanthin), due to β-carotene providing two units of retinol (vitamin A) upon oxidative cleavage in human and animal systems, whereas β-cryptoxanthin provides only one unit. Certain *cis* isomers of β-carotene, namely 9-*cis*-β-carotene and 13-*cis*-β-carotene, have also been of specific interest (to be quantified individually) for human nutrition given that their concentration may change during cooking, and they may have different bioavailability (compared to all-*trans*-β-carotene) during the digestion process (Bohn et al. 2019). Lutein and zeaxanthin are not provitamin A carotenoids, but are nonetheless crucial to human health, specifically eyesight and eye development (Krinsky et al. 2003; Bernstein and Arunkumar 2021). Zeaxanthin is the most abundant grain carotenoid in the population under study herein (Suwarno et al. 2015).

Genomic prediction (GP) methods use genetic markers located throughout the genome to estimate breeding values for phenotypic traits (Crossa et al. 2017). GP can be a valuable tool to accelerate genetic gain in a breeding program by increasing the accuracy of selections and/or reducing cycle time (Crossa et al. 2017). In their review of the implementation of GP in CIMMYT maize and wheat breeding programs, Crossa et al. (2014) described two primary applications: predicting breeding values of individuals for rapid cycling, and predicting genotypic (including epistatic) values of advanced breeding lines; the present study will focus on the former, given the structure of the CAM panel involved herein.

Genomic prediction is typically deployed in one or multiple of four scenarios: I. tested lines are predicted in tested environments; II. tested lines are predicted in untested environments; III. untested lines are predicted in tested environments; IV. untested lines are predicted in untested environments. GP allows the researcher to leverage the known associations between genetic markers and the phenotype in lines from II-IV to make informed predictions. Within-environment prediction is a scenario in which the model is trained on lines grown in a particular environment (location and year) and used to predict the performance of lines grown in that same environment. In between-environment prediction, a model that has been trained on lines grown in one environment is used to predict breeding values for grain carotenoid traits of lines that were grown in a different geographic location and/or year than the lines included in the training set. Predicting the breeding values for one or more phenotypes of interest of lines in an unknown environment using GP could be expected to work best for traits that are highly heritable in the respective target population of environments because the environment has less of an impact on the phenotype.

GP strategies were previously found to exhibit high prediction accuracy for a similar set of grain carotenoid traits (Owens et al. 2014) in 201 inbred lines with yellow to orange grain from a maize diversity panel (Flint-Garcia et al. 2005). That panel consisted primarily of lines with temperate adaptation, with only approximately one-quarter of the lines having tropical/subtropical adaptation (both in the original set of 302 lines and in the 201-line subset analyzed in Owens et al. 2014), and was grown in Indiana, in the Midwest region of the U.S. (Owens et al. 2014). To assess the utility of GP within tropical/subtropical biofortification breeding programs in which MAS has also been deployed, it is important to test GP regression methods and marker sets in a tropical/subtropical maize panel that was grown in the corresponding target environments. These marker sets included *crtRB1*, as well as *lcyE* (*lycopene ε-cyclase* which encodes the enzyme at the branchpoint of the core carotenoid pathway) that have known alleles that are relevant for biofortification in the CIMMYT program (Yan et al. 2010; Babu et al. 2013; Suwarno et al. 2015) and that have been identified in genome-wide association studies for grain provitamin A (and other carotenoid) traits in maize (Harjes et al. 2008; Yan et al. 2010; Owens et al. 2014; Suwarno et al. 2015; Azmach et al. 2018; Baseggio et al. 2020). We also tested a larger set of 13 genes that were identified for grain carotenoid traits in the U.S. maize nested association mapping panel (Diepenbrock et al. 2021; LaPorte et al. 2022; Table S1), for which approximately half of the parents were tropical in adaptation. The results from a GP analysis of the CAM panel could help increase genetic gain for maize grain carotenoid traits in this breeding program. The high heritabilities of the carotenoid traits in this panel (as reported in Suwarno et al. 2015) make them good candidates for GP given that heritability represents the proportion of phenotypic variance that is explained by genetic factors. Implementation of GP strategies has been recently explored to support the biofortification of vitamin E (Tibbs-Cortes et al. 2022) and zinc-related (Guo et al. 2020) traits in maize, iron and zinc (among other health-beneficial mineral nutrients) in wheat, and zinc in rice (Rakotondramanana et al. 2022).

Three types of regression have been extensively used to train GP models (Usai et al. 2009; Heslot et al. 2012; Azodi et al. 2019): Ridge Regression-Best Linear Unbiased Prediction (RR-BLUP; Meuwissen et al. 2001; Endelman 2011), Least Absolute Shrinkage and Selection Operator (LASSO; Tibshirani 1996), and Elastic Net (Zou and Hastie 2005). These methods are regularization techniques, meaning they penalize model complexity and differ based on their penalization strategies (Ogutu et al. 2012). In brief, the penalization strategy for Ridge Regression (L2 regularization) is the sum of squares of the coefficients of the model. By contrast, LASSO (L1 regularization) uses the absolute value of the coefficients and will zero coefficients for a sparse solution. Elastic Net is an intermediate method that utilizes both L1 and L2 regularization, modulated by a hyperparameter, α (Zou and Hastie 2005). A Reproducing Kernel Hilbert Space (RKHS)-based approach has been developed for GP to take advantage of semi-supervised learning based on a projection into a Hilbert space (which is a mathematical construct that allows for such a projection) to model non-linear relationships between predictors and traits more accurately (Gianola et al. 2006; Campos et al. 2010; Montesinos López et al. 2022). Kernel approaches, including RKHS, theoretically will have higher prediction accuracy for traits with a more complex genetic architecture, that cannot be described by a linear model, underpinning the phenotype (Montesinos López et al. 2022).

Implementation of GP relies on similar data sets as those used for genetic mapping—i.e., genotypic and phenotypic data. As such, being able to conduct GP in the same analytical workflow as genetic mapping and other quantitative genetics analyses could offer benefits from a computational efficiency perspective and alleviate barriers to entry for GP approaches that otherwise require custom scripting (and, for some implementations, high-performance computing). A software package called Trait Analysis by aSSociation, Evolution and Linkage (TASSEL; Bradbury et al. 2007) is widely used among the plant breeding/genetics community for quantitative genetics analyses. TASSEL is well documented and maintained, can be operated in a graphical user interface (which is also helpful for alleviating barriers to entry), and is configured to accept multiple common formats of genotypic and phenotypic data.

This study investigates the prediction accuracies of several GP models to inform the implementation of genomic prediction/selection (GP/GS) in breeding efforts for carotenoid-dense tropical and subtropical maize. We compared several GP methods to determine the most accurate method for predicting grain carotenoid traits within environments, including RR-BLUP, the TASSEL Genomic Selection plugin implementation of GBLUP, LASSO, Elastic Net, and RKHS approaches. Additionally, we tested the efficacy of RR-BLUP to predict between environments. Furthermore, we compared the accuracy of models trained on genome-wide marker data vs. markers proximal to two or 13 genes of known relevance to grain carotenoid biosynthesis within and between environments. Finally, we describe the performance of a Genomic Selection plugin that has been implemented in TASSEL for use by researchers and test the prediction accuracy and computational efficiency of this plugin, including in comparison with other methods.

## Methods

### Maize Germplasm

The design of the CAM panel was described in Suwarno et al. (2015). Briefly, inbred lines were selected at the initial stages of the CIMMYT-HarvestPlus maize biofortification breeding program, more than a decade ago, for inclusion in the panel. The CAM panel includes 380 lines, with the majority being tropical or subtropical lines (47% each) and 3% of lines adapted to temperate conditions (Suwarno et al. 2015). In addition to the CAM panel, 65 other CIMMYT lines were quantified in the same environments and were included in GP. These lines were distributed throughout the population in terms of genomic relatedness (Figure S1). Of the included maize lines, 10 lines in this study include the beneficial alleles of one gene associated with high provitamin A (*crtRB1*). The pedigree of these lines was reported by Suwarno et al. (2015). These lines were: CIM-SYN-386-34, CIM-SYN-387-35, CIM-SYN-389-37, CIM-SYN-390-38, CIM-SYN-391-39, CIM-SYN-392-40, CIM-SYN-393-41, CIM-SYN-394-42, CIM-SYN-415-63, and CIM-SYN-416-64. The CAM panel was grown in Agua Fría (AF) in the state of Puebla, Mexico, in the years 2012 and 2013, and Tlaltizapan (TL) in the state of Morelos, Mexico, in the years 2010 and 2011, resulting in four environments hereafter referred to as the combination of location and year: TL10, TL11, AF12, and AF13 (Suwarno et al. 2015; Figure S4). Tlaltizapan (which is considered mid-altitude subtropical) is located 268 km south and west of Agua Fría (which is considered tropical). Tlaltizapan had, at the time of experimentation, a slightly warmer (+1.5 °C), drier (−360 mm/year) climate at a higher elevation (+835 masl) than Agua Fría (Suwarno et al. 2015). Tlaltizapan and Agua Fría are in Aw (class 3) and Am (class 2), respectively, according to the Köppen-Geiger classification (Beck et al. 2018; Figure S4).

### Phenotypic Datasets

Lines from the four distinct environments (AF12, AF13, TL10, TL11) were phenotyped for the following traits: lutein, zeaxanthin, β-carotene, β-cryptoxanthin, and provitamin A (defined as the concentration of β-carotene + 0.5(β-cryptoxanthin)) as described in Suwarno et al. (2015). In the Agua Fría environments, lines were also phenotyped for two isomers of β-carotene: 9-*cis*-β-carotene and 13-*cis*-β-carotene. Briefly, two to six plants per plot were self-pollinated, and random samples of 50 seeds from each accession were ground and quantified for carotenoid analysis following the CIMMYT laboratory procedure (Galicia et al. 2009; Palacios-Rojas et al. 2017). In samples collected from TL10 and TL11, concentrations of individual carotenoid compounds were quantified using HPLC and in Agua Fría, they were quantified using UPLC (Suwarno et al. 2015). The results were reported as μg g^-1^ dry kernel weight (Suwarno et al. 2015; Galicia et al. 2009).

### Data Preparation

Genotypic data for each accession was generated as described by Suwarno et al. (2015). Briefly, genotyping by sequencing (GBS) (955,120 SNPs) was used for these lines, as generated by the Institute for Genomic Diversity, Cornell University, Ithaca, NY, USA (Suwarno et al. 2015). Filtering for minor allele frequencies (MAF) was not conducted before GP because monomorphic markers can be informative for prediction. The genotypic data, in the form of a matrix of SNP markers (including monomorphic markers), was numericalized using the HapMap() function from Genome Association and Prediction Integrated Tool (GAPIT) version 3 in R (R Core Team 2021; Wang and Zhang 2021) for all methods except for the TASSEL plugin, for which a kinship matrix was calculated within TASSEL from the input genotypic data set as described below. The phenotypic data for each line and trait was matched to the genotype using the accession identifier.

### Model Types

The model types used in the model fitting procedure described above include RR-BLUP, Least Absolute Shrinkage and Selection Operator (LASSO), Elastic Net (EN), and Reproducing Kernel Hilbert Spaces (RKHS). RR-BLUP was completed using the rrBLUP package (Endelman 2011) function kinship.BLUP(). This function carries out both the training and prediction steps. The median and standard deviation (SD) of prediction accuracy across the five folds was calculated and reported. LASSO was fit using the GLMNet package (Friedman et al. 2010) function cv.glmnet(), with α = 1 and by tuning the lambda hyperparameter (the shrinkage parameter) by identifying the value with the lowest mean cross-validated error (referenced in the documentation as ‘cvm’) from a set of 250 potential values for each fold and iteration, using the nlambda argument (Friedman et al. 2010). Predictions were calculated using the GLMNet library function predict(). Within each cross-validation model fitting instance, Elastic Net was conducted nine times using the GLMNet() function from the GLMNet package (Friedman et al. 2010), with a different α value each time. The nine α values (the weight of L1 versus L2 regularization penalties) tested ranged from 0.1 through 0.9 with steps of 0.1. Note that an α value of 0 is equivalent to the LASSO method, and an α value equal to 1 corresponds to RR-BLUP in this function. The lambda (shrinkage) parameter was selected from a set of 250 values for each α value of Elastic Net separately, in the same manner as described above for LASSO.

The BGLR (Bayesian Generalized Linear Regression) function from the BGLR package was used to train and fit a Reproducing Kernel Hilbert Space (RKHS) model (Perez and Campos 2014). The Gaussian kernel was used as the covariance matrix (Kernel ‘K’), calculated using a bandwidth parameter of 1 and a distance matrix, D, containing the Euclidean distance between the training and test set of marker genotypes (a square matrix with the dimensions of the number of lines). We ran 6000 iterations per fit model with a burn-in size of 1000 (based on hyperparameter tuning with smaller burn-in sizes). The bglr() function was used to fit the model and generate the predicted values.

The GBLUP method was conducted using the TASSEL GUI built-in implementation (Bradbury et al. 2007). After genotypic data was loaded, a kinship matrix was created using TASSEL. Genomic prediction was conducted by selecting the phenotypic file and kinship matrix. Five-fold cross-validation was repeated in 20 iterations for each trait and environment. While all other methods (Ridge, EN, LASSO, RKHS) used the same line assignments for the training and test sets, TASSEL did not, as the plugin conducts random fold assignments unless given an incomplete phenotype vector (in which values in the desired test set have been masked). The RKHS, LASSO, and RR-BLUP methods were run with the same numericalization strategy and training and test splits (within a given iteration), and are thereby a direct comparison within a given trait. Running TASSEL on the same train-test-split assignments as the other methods was performed for a subset of iterations to determine the extent to which RR-BLUP predicted values were similar to those generated by TASSEL. RR-BLUP, RKHS, LASSO, and EN were run on a High-Performance Computing (HPC) cluster, with 250G of memory allocated to a parallelized script. The TASSEL GUI and the two-gene and 13-gene R scripts were all run on the desktop. The TASSEL software was run with a maximum heap of −Xmx 100g.

We also compared the use of subsets of known carotenoid-related genes in prediction to whole-genome prediction. Two-gene and 13-gene approaches were conducted with RR-BLUP, both within and between environments, using the markers within 250 kilobases (kb) of genes that are related to grain carotenoid traits (Table S1); these markers were extracted using a custom R script. The locations of the genes used in the two-gene and 13-gene approaches were determined using MaizeGDB (Woodhouse et al. 2021; Table S1). The two-gene approach included all markers within ± 250 kb of *lcyE* and *crtRB1* (1,540 markers). The 13-gene approach included markers within ± 250 kb of 13 identified carotenoid-related genes (11,143 markers) (Diepenbrock et al. 2021; LaPorte et al. 2022; Table S1), which included *lcyE* and *crtRB1*.

Between-environment analyses were conducted using RR-BLUP. Every pairwise combination of environments (referenced here as environments E1 and E2) were evaluated as training and test sets for prediction. Five-fold cross validation was conducted for each pair of environments E1 and E2, such that both within- and between-environment prediction were conducted in an identical framework. Namely, for a given fold of cross-validation, 1) 80% of E1 was used as the training set, and the remaining 20% of E1 was used as the test set; 2) 80% of E2 was used as the training set, and the remaining 20% of E2 was used as the test set; 3) 80% of E1 was used as the training set, and the remaining 20% of E2 was used as the test set; and finally, 4) 80% of E2 was used as the training set, and the remaining 20% of E1 was used as the test set.

Genomic prediction (GP) was conducted using a five-fold cross-validation approach with 20 iterations. As a result, each model was run 100 times. For each cross-validation instance, the accessions were divided into randomized training and testing sets, with ~80% of the lines used for training and the remaining ~20% withheld for testing model performance. Each GP model was trained using genomic and phenotypic data in the training set, and the resulting model was then fit to the test set to predict their breeding values based on genomic information. Accuracy for every model type was calculated as the Pearson correlation (*r*) between the predicted values for the test set and the actual observed values in the test set. The median prediction accuracy of each iteration was calculated and reported. The 20-iteration, five-fold cross-validation process was repeated with each model-fitting method (RR-BLUP, EN, LASSO, and RKHS) for each of the carotenoid traits within each environment, using the same lines in the training and test sets for each method. For within-environment GP, the training and test sets were grown in the same environment (AF12, AF13, TL10, TL11). The five-fold cross-validation process was also conducted with RR-BLUP between each environment for each of the carotenoid traits (β-carotene, β-cryptoxanthin, lutein, zeaxanthin, and provitamin A in all environments, as well as 9-*cis*-β-carotene and 13-*cis*-β-carotene in the Agua Fría environments).

Median prediction accuracies were depicted in figures as a measure of central tendency for accuracy because they are less affected by outliers than means. Plots for all models were visualized using ggplot2 and color palettes from Viridis and base R (Wickham 2016; R Core Team 2021; Garnier et al. 2023). Models are systematically referred to as “similar to” and “different from” one another based on the differences in median prediction accuracy between all iterations for a given model type, trait, and environment. Namely, “similar to” refers to models where the median accuracy between all iterations of one method was within the interquartile range (IQR) of prediction accuracies of another method. Models being “better than” (or “worse than”) one another is defined as when the median for the “better” (or worse) method is higher (or lower) than the median of the second method and not within the IQR of that method.

## Results

### Phenotypic correlations

The phenotypic values were positively correlated for most traits and in most environments (Figure 1). Zeaxanthin levels in all environments, and lutein in the Agua Fría environments were highly positively correlated. A high positive correlation was observed between provitamin A and β-carotene in all environments. A negative correlation was observed between lutein and each of provitamin A and β-carotene in the Tlaltizapan environments. β-cryptoxanthin showed a slightly higher correlation with provitamin A within Agua Fría environments than within the Tlaltizapan environments.

**Figure 1.**
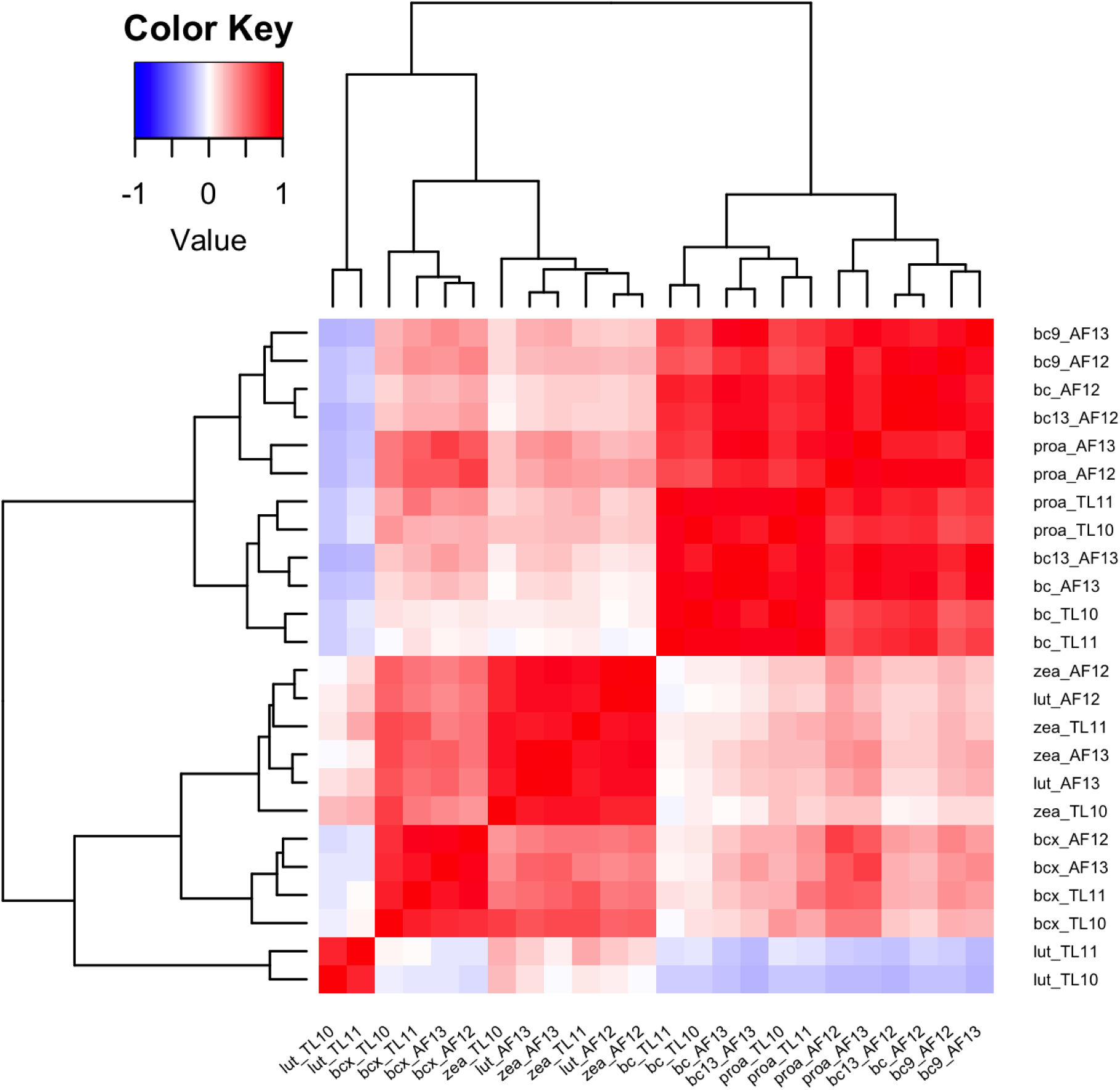
Correlations between phenotypic values for each trait and environment. The trait-environment pairs are listed as the abbreviation for the trait (lut, lutein; zea, zeaxanthin; bcx, β-cryptoxanthin; bc, β-carotene; proa, provitamin A) and the environment (Agua Fría 2012, Agua Fría 2013, Tlaltizapan 2010, and Tlaltizapan 2011 [AF12, AF13, TL10, and TL11, respectively]). The heatmap is arranged in clustered groups denoted with the line diagram. Red corresponds to a positive correlation, and blue corresponds to a negative correlation. The darker the color, the higher the absolute value of the correlation

### Within-environment prediction

First, we compared the prediction accuracy of RR-BLUP (α = 0), Elastic Net (α = 0.1 to 0.9), and LASSO (α = 1) (Figure 2). For most traits and in most environments, RR-BLUP had the highest prediction accuracies, although in some cases, the prediction accuracy was similar to that of one or several EN models. Specifically, RR-BLUP had better prediction accuracy than all other methods for provitamin A (in TL11, AF13, and AF12), zeaxanthin (in TL10 and AF13), β-cryptoxanthin (in TL10 and AF13), 9-*cis*-β-carotene (in TL11 and AF13), β-carotene (in TL10), and 13-*cis*-β-carotene (in AF12). For most trait-environment combinations, LASSO performed similarly to the EN models except in the case of three traits in TL11. Specifically, LASSO performed worse than all EN methods and RR-BLUP for zeaxanthin in TL11, and LASSO and RR-BLUP performed better than all of the EN models for β-carotene and provitamin A in TL11. High prediction accuracies were observed for β-carotene and provitamin A in TL10 for all of RR-BLUP, EN, and LASSO, with the medians of all models higher than 0.75 for β-carotene and ranging from 0.71 to 0.75 for provitamin A. However, β-carotene and (to a lesser extent) provitamin A also generally had larger ranges of median prediction accuracy (represented by the height of the violins; Figure 2) than the other traits in each environment. For this within-environment prediction experiment, RR-BLUP took the least computational time compared to EN and LASSO (which took approximately 4 times as long).

**Figure 2.**
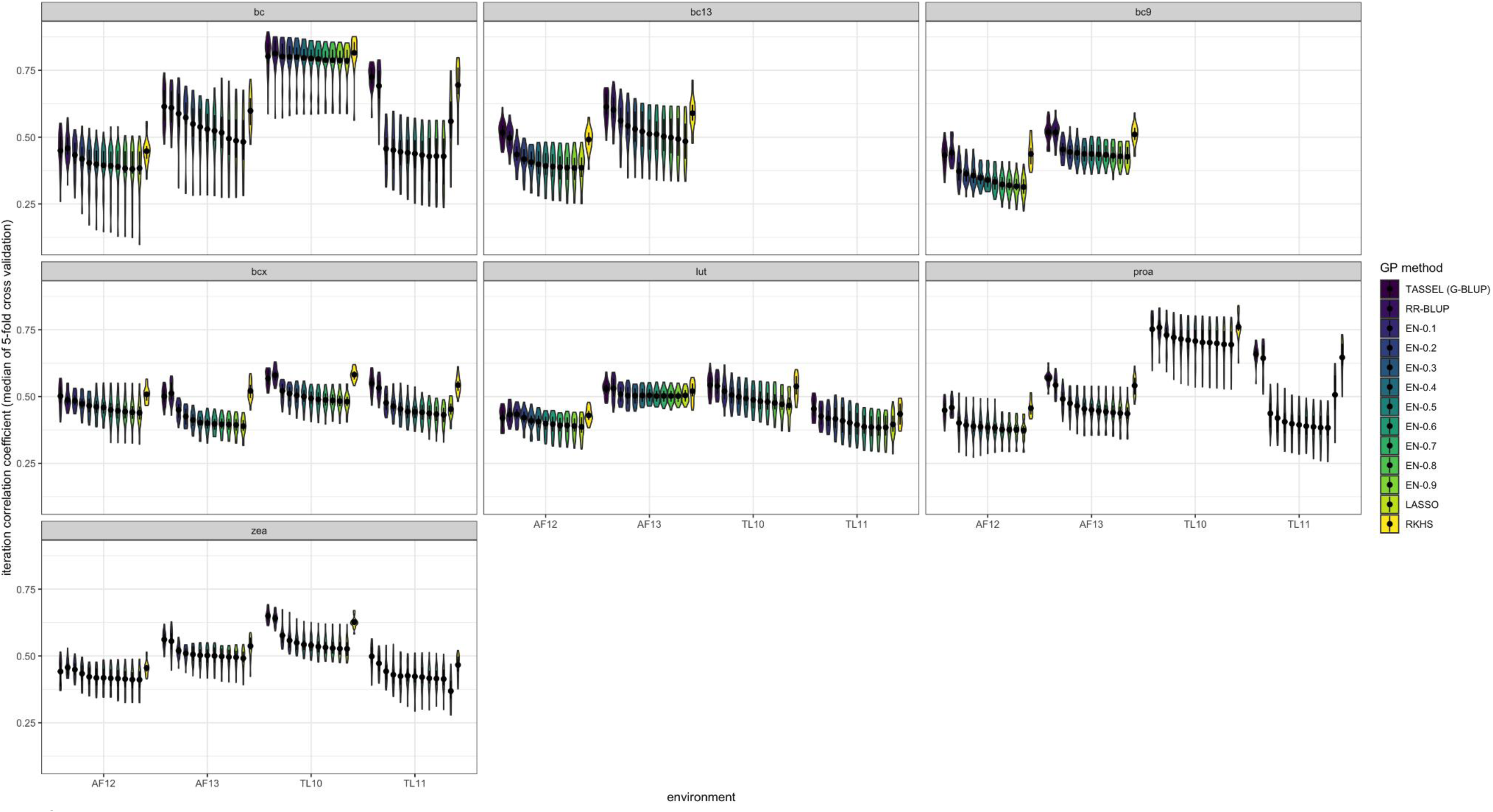
Within-environment prediction accuracies for five regression types (TASSEL, Ridge, Elastic Net, LASSO and Reproducing Kernel Hilbert Space) for each trait-environment combination. Ridge Regression Best Linear Unbiased Prediction = RR-BLUP; LASSO = Least Absolute Shrinkage and Selection Operator; EN = Elastic Net; RKHS = Reproducing Kernel Hilbert Space; TASSEL = the TASSEL GUI Genomic Selection Plugin implementation of G-BLUP. For EN, the numeric value of 0.1 to 0.9 (in steps of 0.1) indicates the ɑ value used in EN. Trait abbreviations: bc, β-carotene; bc9, 9-cis-β-carotene; bc13, 13-cis-β-carotene; bcx, β-cryptoxanthin; lut, lutein; proa, provitamin A; zea, zeaxanthin. The median for each violin is represented by a black dot. The interquartile range (Q1-Q3) of each set of iterations is represented by a black vertical line through the center of each violin.

The TASSEL GS plugin, RR-BLUP, and RKHS had similar accuracies for all traits and environments (Figure 2). In all trait-environment combinations, these three methods performed better than LASSO (which exhibited median prediction accuracies of 0.10 to 0.59). Although there were some variations in the median accuracy, none of the median accuracies for these three superior-performing methods were outside of the IQR of one another for a given trait-environment combination (0.32 to 0.88 for RR-BLUP, 0.26 to 0.89 for TASSEL, and 0.34 to 0.88 for RKHS). For RR-BLUP and TASSEL, the prediction accuracy was highest for β-carotene (highest median values of 0.831 and 0.821, respectively, for any trait-environment combination). Although the RKHS method performed as well as RR-BLUP and TASSEL in all instances, it took longer to run (approximately 6 times as long).

We plotted the median prediction accuracy for each iteration of the RR-BLUP within-environment prediction experiment versus the SD of prediction accuracy across folds within that iteration (Figure S2). No clear trend was observed between these two metrics for most traits and most environments. For certain traits and environments—for example, β-carotene in all environments—iterations with higher SD tended to have lower prediction accuracy. The raw phenotypic values were also examined to check for any relationship with SD of prediction accuracy within iterations; the range of phenotypic values was similar between the environments (Figure S3). The ten biofortified lines in the panel under study herein had an allele of *crtRB1* that was previously found to be favorable for provitamin A (Yan et al. 2010; Babu et al. 2013; Suwarno et al. 2015). The lines that possessed these alleles indeed had among the highest levels of β-carotene and provitamin A (Figure S3). For the other traits, notably including β-cryptoxanthin, the phenotypic values were located throughout the population distribution.

### Between-environment prediction

RR-BLUP was performed between environments to test how accurately the data from one environment could be used as a training set to predict values for a different environment. Between-environment predictions performed approximately as well as within-environment predictions for the lowest-prediction accuracy environment (Figure 3). Notable exceptions include predictions for lutein in instances in which data from Agua Fría (AF) are being used to predict in Tlaltizapan (TL) and vice versa; in each of those cases, the prediction accuracy for lutein was near zero. The within-environment prediction accuracies for TL10 for β-carotene and provitamin A were higher, and narrower in range (indicated by the height of the violin; Figure 3), compared to the between-environment prediction accuracies for TL10 when AF13 was used as the training set. When different years were compared in the same location (between AF12 and AF13, or between TL10 and TL11), the accuracy was similar to those of the across-location models. In some cases, there was a marginally higher prediction accuracy when an environment was used as the test rather than the training set. For example, higher prediction accuracy was observed for provitamin A when AF12 was used as the training set (0.531) and TL11 as the test set rather than the opposite scenario (0.456).

**Figure 3.**
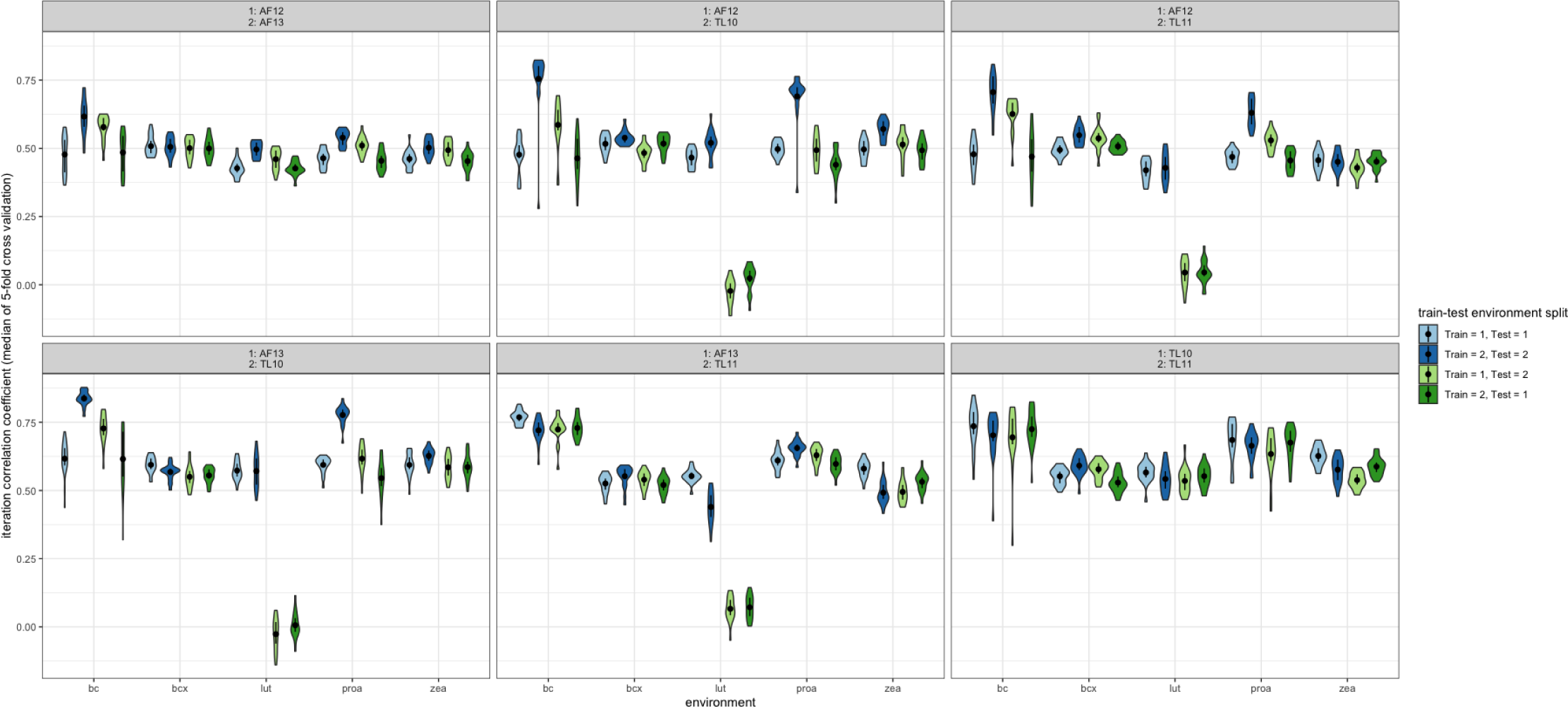
Between-environment prediction accuracies using RR-BLUP for all traits, within every pairwise combination of environments. Trait abbreviations: bc, β-carotene; bc9, 9-cis-β-carotene, bc13, 13-cis-β-carotene; bcx, β-cryptoxanthin; lut, lutein; proa, provitamin A; zea, zeaxanthin

### Testing of marker sets

Next, we investigated if including fewer markers in RR-BLUP could have comparable accuracy to the best-performing whole-genome prediction methods (RR-BLUP and TASSEL). For this purpose, we incorporated two additional approaches, referred to as the two-gene and 13-gene approaches (Table S1), that utilized a smaller subset of markers as the only predictors in the model. The two-gene and 13-gene approaches (using RR-BLUP) both performed with lower prediction accuracy than genome-wide approaches that used RR-BLUP or the TASSEL implementation of GBLUP. The 13-gene approach performed much better than the two-gene approach for some trait-environment combinations: notably, lutein in the AF environments and zeaxanthin in all environments. The 13-gene approach performed better than LASSO (except for similar performance in AF12), which in turn performed better than the two-gene approach, for β-cryptoxanthin. For β-carotene and provitamin A, the two-gene and 13-gene approaches had the same prediction accuracy. For 13-*cis*-β-carotene and 9-*cis*-β-carotene, the two-gene method performed better than the 13-gene method. The two- and 13-gene approaches maintained similar prediction accuracies between vs. within environments for all traits except for lutein, for which accuracies were near zero between locations (particularly for the two-gene approach; Figures S5-S6).

## Discussion

Genomic prediction is the central mathematical exercise in Genomic Selection. A key application of these models is the prediction of germplasm performance in tested and untested environments (Heslot et al. 2012, 2015; Bhat et al. 2016; Azodi et al. 2019). In this study, we predicted tested lines in tested environments and those same lines in “untested” environments (for which phenotypes had been masked) through between-environment prediction. Knowing the accuracy of a model within and/or between environments is helpful for breeders who wish to use this model to predict the performance of tested and/or untested lines.

### Predicting with models that use fewer markers

RR-BLUP consistently exhibited a higher prediction accuracy than LASSO. These results suggest that, in this dataset, penalizing model complexity and utilizing a smaller set of predictors was detrimental to model accuracy, even for these grain carotenoid traits, which are thought to be relatively oligogenic in architecture. Incorporating more markers appears to be beneficial for model accuracy within the model configurations tested within this study. This finding is corroborated by the results in Figure 4, where the two- and 13-gene approaches had a lower prediction accuracy than the RR-BLUP and TASSEL plugin (GBLUP) approaches that included genome-wide markers. LASSO, which also tested a smaller subset of markers (that the model identifies through hyperparameter optimization) but which uses no prior information about the markers in the selection of the subset, performed worse than the 13-gene method (but better than the two-gene method) for β-cryptoxanthin in three of four environments, and otherwise performed similarly to the 13-gene method (Figure 4). In this study, whole-genome prediction methods were more accurate than methods that used marker subsets. The two-gene approach was tested given the use of *lcyE* and *crtRB1* in MAS efforts specifically for provitamin A, whereas the 13-gene approach was tested to determine the accuracy of a model focused on a broader set of *a priori* genes for grain carotenoid traits. Based on the two-gene approach exhibiting worse prediction accuracies than the 13-gene approach (and approaches using genome-wide markers) for zeaxanthin within each of the four environments, we particularly recommend using a marker set that is less targeted than the two-gene subset for prediction of or selection upon zeaxanthin.

**Figure 4.**
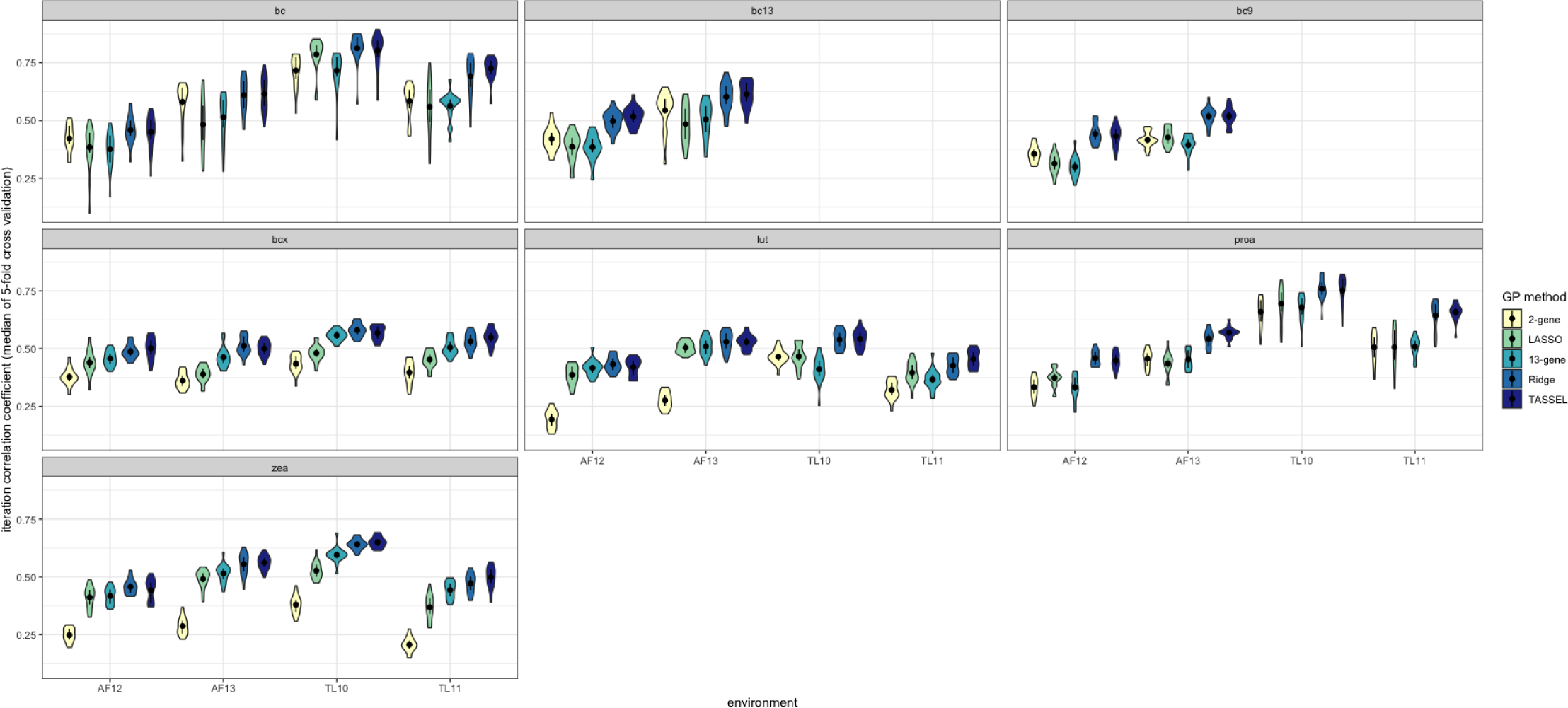
Comparison of prediction accuracies for three model types (RR-BLUP, TASSEL (GBLUP), and LASSO) using genome-wide markers or RR-BLUP using markers proximal to two or 13 genes. Trait abbreviations: bc, β-carotene; bc9, 9-*cis*-β-carotene, bc13, 13-*cis*-β-carotene; bcx, β-cryptoxanthin; lut, lutein; proa, provitamin A; zea, zeaxanthin

### Between-environment prediction

RR-BLUP was used for between-environment prediction because it had the highest accuracy and computational efficiency of all regression methods (LASSO, EN, RKHS, Ridge) tested in the within-environment experiment (Figures 1, 2, 3). The between-environment models maintained prediction accuracy between locations and years (with the exception of lutein between locations), which is concordant with the high heritability of the tested traits, and suggests that the results of these predictions could be beneficial for breeders wishing to develop carotenoid-dense varieties for untested environments (at least within a similar climatic zone). Breeding of provitamin A-dense maize varieties is also taking place by teams at CIMMYT, IITA and other institutions in southern and western Africa, among other geographies. Testing prediction accuracy in multi-environment trials across CIMMYT regional stations (namely between Mexico and sub-Saharan Africa) would be informative in optimizing collaborative breeding activities across those stations vs. station-specific activities focused on local/subregional adaptation. Namely, GP (and estimates of heritability) across mega-environments would further inform the extent to which grain carotenoid traits could be treated as largely genetic parameters or whether they would need to be treated more akin to agronomic traits (for which breeding for local/subregional adaptation has been critical within provitamin A biofortification efforts; Manjeru et al. 2019). The two locations examined in the present study had the same Köppen-Geiger classification (Figure S4), which may impact the ability to extrapolate beyond this environmental class.

### Between-environment prediction with fewer markers

Prediction accuracies were maintained between vs. within environments for the two-and 13-gene approaches, indicating that markers proximal to these genes were able to predict consistently despite environmental effects. Owens et al. (2014) tested the use of random sets of eight genes (selected from the whole genome or from a set of 58 *a priori* genes for maize grain carotenoids) in prediction and found that a non-random set of eight major-effect *a priori* genes (which were included in the 13 genes tested herein) exhibited improved performance compared to both types of random sets. That finding suggests that the non-negligible prediction accuracies of the two- and 13-gene approaches observed herein (particularly within environments, and for provitamin A traits between environments) were not solely due to gene (and corresponding marker) subsets of those sizes simply managing to sufficiently capture genomic relationships—rather, that gene identity (and namely relevance to the target traits) is important. Nonetheless, given that genome-wide markers exhibited high predictive abilities in this study, and given that GP/GS is deployed at CIMMYT for other traits (such that assaying of genome-wide markers could be beneficial for other traits as well) and such assays are increasingly low-cost, it seems that use of genome-wide markers in GS for grain carotenoid traits would be robust (with moderate to high prediction accuracies observed in this study both within and between environments; with the exception of lutein between locations, which showed poor prediction accuracies for all marker sets) and still resource-efficient.

### Considerations for lutein quantification and prediction in carotenoid biofortification programs

In this study, trait values for lutein in TL10 and TL11 showed slight negative correlations with β-carotene and provitamin A and slight to negligible correlations with β-cryptoxanthin and zeaxanthin in all environments (Figure 1). While trait values for lutein in AF12 and 13 had slight to negligible correlations with β-carotene, they exhibited a slight positive correlation with provitamin A and substantially higher positive correlations (than those observed for lutein in the TL environments) with β-cryptoxanthin (median 0.45, SD 0.06) and zeaxanthin (median 0.81, SD 0.08) in all environments. These relationships would be important to monitor if seeking to increase lutein (produced in the ɑ-branch of the carotenoid pathway) alongside other carotenoid traits (namely those produced in the β-branch, including β-carotene, β-cryptoxanthin, and zeaxanthin)—particularly if selecting upon *lcyE,* which encodes the enzymatic step at the pathway branchpoint.

Interestingly, these trait relationships were consistent across years within locations. One potential explanation for the differential relationships observed between locations could be that some environmental (or repeatable GxE) factor in TL vs. AF made for weaker vs. stronger relationships of lutein with other traits (namely β-cryptoxanthin and zeaxanthin) in the two locations. While we do not exclude this possibility, it was also noted in Suwarno et al. (2015) that the HPLC method used for carotenoid quantification in TL10 and 11 allowed for better separation of lutein and zeaxanthin on the resulting chromatograms than the UPLC method used for AF12 (and the same UPLC protocol was used for AF13). Notably, the accuracy of GP models for lutein was poor when predicting between TL and AF environments in either direction. That finding could be due to lack of concordance between lutein as quantified via HPLC vs. UPLC, as instrument platform was confounded with location. The substantially higher correlations observed between lutein and zeaxanthin in AF (compared to TL) could be artificially high due to that inferior separation, as a larger peak in the relevant range of retention times could have resulted in higher quantified levels of both compounds. Taken together, we would recommend prioritizing the prediction of values acquired through optimal analytical methods for all analytes and gathered using the same instrument platforms.

### Suitability of the TASSEL GS plugin for GP

The TASSEL GS plugin is accessed through the TASSEL GUI; the results are generated quickly (in approximately 30 seconds), and all traits can be run simultaneously. This plugin could be helpful for researchers needing to perform GP without HPC resources or who wish to test the model as part of their quantitative genetics analytical workflows (e.g., if genetic mapping is already being conducted in TASSEL) without building up dedicated custom script bases, which can represent a barrier to entry. This plugin is a user-friendly resource that is hosted within the TASSEL platform and community, with thorough documentation and troubleshooting resources. The TASSEL GS plugin can also be used for species other than maize (and TASSEL as a platform has been used widely across species). Although the GS plugin has been available in TASSEL since 2015, its accuracy compared to other GP methods has not yet been compared in the literature until the present study. The high accuracy of the plugin, on par with that of RR-BLUP, demonstrates that TASSEL provides a highly accurate method for GP that could aid breeders of many different crops in GP/GS. This is to be expected given that RR-BLUP and G BLUP (the latter is used by TASSEL) have been demonstrated to be mathematically equivalent (Meuwissen et al. 2001; Habier et al. 2007; Clark and Van Der Werf 2013). We would also refer readers to rTASSEL (Monier et al. 2022), which offers many of the same utilities of TASSEL (including GP) in an R environment. We anticipate that both the TASSEL GUI and rTASSEL implementations could have utility for partially overlapping user bases, with the overall outcome of increased accessibility to GP methods and increased continuity with other analytical workflows routinely carried out by plant breeders/geneticists. Finally, the kin.blup() function has superceded kinship.blup() in the rrBLUP package (Endelman et al. 2011); while both are valid from a linear algebra perspective (and are wrappers to the mixed.solve() function in the same package), kin.blup() offers the advantage of the user not needing to specify design matrices.

### Considerations for implementing GS in biofortification programs

Ten lines in this study were biofortified for β-carotene and had among the highest grain concentrations of β-carotene in the panel (Figure S3; Suwarno et al. 2015). Twenty iterations were conducted per method per experiment, largely to mitigate the placement of a small number of lines within a given randomized fold assignment influencing the overall median prediction accuracy (as calculated across iterations). The iterations with the highest prediction accuracy generally had low SDs between the different cross-validation folds within that iteration. The median prediction accuracy in most traits and environments had either no linear trend or a negative linear trend when plotted against SD, a metric for variation or dispersion, within model iterations (Figure S2).

Provitamin A is a ‘derived’ trait in the sense that it was calculated as the concentration of β-carotene plus half of the concentration of β-cryptoxanthin. Given that β-cryptoxanthin is immediately downstream of β-carotene within the biosynthetic pathway, conducting GP for provitamin A itself—as a trait that is of direct interest for human nutrition—is important to biofortification efforts, and both accounts for and masks the biological complexity underlying provitamin A concentrations. *crtRB1* encodes an enzyme that catalyzes two consecutive steps in the carotenoid biosynthetic pathway: β-carotene to β-cryptoxanthin, and β-cryptoxanthin to zeaxanthin. This gene was found to exhibit negative pleiotropy between β-carotene and each of β-cryptoxanthin and zeaxanthin in the U.S. maize NAM panel (Diepenbrock et al. 2021). The 10 lines with a favorable allele of *crtRB1* that were included in this study had among the highest levels of β-carotene, but their levels of β-cryptoxanthin were distributed throughout those of the rest of the population (Figure S3). Use of GP, whether using genome-wide markers or markers proximal to a larger set of genes, could help in optimization of provitamin A concentrations as the sum of the two compound concentrations without solely operating upon their direct substrate-to-product relationship, which is itself upstream of another health-beneficial carotenoid: zeaxanthin.

### Future directions and recommendations

Next steps for this work include the testing of grain carotenoid concentrations (and GP models predicting them) across mega-environments. Notably, we would recommend ensuring that the same instrument platform was used for carotenoid quantification when conducting single-trait predictions across environments, particularly for lutein. If UPLC methods that were specifically optimized for provitamin A carotenoids are being used, multi-trait GP models (based on levels of provitamin A compounds) may have higher accuracies for lutein (and perhaps secondarily, zeaxanthin) than single-trait models that train and predict breeding values for those non-provitamin A compounds on data from different analytical platforms. However, such approaches would be limited by the extent of the genetic correlations between levels of provitamin A and non-provitamin A compounds (which could also change as one or both trait sets are being selected upon) and would need to be tested empirically.

Another key next step for this work is the use of GP methods in selection. Given that the CAM panel is composed of key inbreds for the breeding program, the recurrent selection scheme proposed by Windhausen et al. (2012) for application of GP in closed populations could be pertinent. Namely, recurrent selection (repeated cycles of selection with intermating of selected individuals, in distinct rather than overlapping rounds; Labroo and Rutkoski 2022) could be used to conduct population improvement for priority traits and select candidate pre-commercial lines out of that population. Key inbreds from outside of the CAM panel could also be incorporated to help maintain genetic variation and integrate complementary favorable genomic regions (Paula et al. 2020). We would recommend using GP methods that penalize or monitor for loss of genetic diversity; Ozimati et al. (2019) did not observe such a loss when carrying out GS for four traits including dry matter content in cassava, though that program prioritized crosses across (rather than within) population clusters based on genomic relationships. The TASSEL GS plugin can be readily tested by interested parties on the TASSEL tutorial data, which are included in the TASSEL installation directory, and/or on the input data sets used in the present study. In conclusion, high accuracies of GP methods for maize grain carotenoid traits herein, even via less computationally intensive methods, and the availability of the TASSEL GS plugin represent the predictive ability of and an opportunity for the use of GP/GS in biofortification alongside other crop improvement efforts.

## Acknowledgments

We gratefully acknowledge Raman Babu and Kevin Pixley for their involvement in the development of the CAM panel and Xuecai Zhang and Daniel Runcie for providing constructive feedback on this manuscript. We also thank the International Maize and Wheat Improvement Center and Harvest Plus for having supported the development of the CAM panel. We also thank Jaime Cuevas Domínguez for his help with the BGLR library. We thank Terry M. Casstevens, Edward S. Buckler, and the rest of the USDA-ARS supported TASSEL developer team for their involvement in developing and maintaining the TASSEL software. We also thank Jorge Santiago Ramírez-Núñez for help with GIS mapping software. This material is based upon work supported by the U.S. Department of Energy, Office of Science, Office of Advanced Scientific Computing Research, Department of Energy Computational Science Graduate Fellowship under Award Number DE-SC0021110.

## Statements and Declarations

### Disclaimer

This report was prepared as an account of work sponsored by an agency of the United States Government. Neither the United States Government nor any agency thereof, nor any of their employees, makes any warranty, express or implied, or assumes any legal liability or responsibility for the accuracy, completeness, or usefulness of any information, apparatus, product, or process disclosed, or represents that its use would not infringe privately owned rights. Reference herein to any specific commercial product, process, or service by trade name, trademark, manufacturer, or otherwise does not necessarily constitute or imply its endorsement, recommendation, or favoring by the United States Government or any agency thereof. The views and opinions of authors expressed herein do not necessarily state or reflect those of the United States Government or any agency thereof.

### Author Contributions

MFL, WBS, JC, NPR and CHD planned and designed the research. MFL, AK, and CHD conducted the analyses. PH, NPR generated a portion of the data used in this study, alongside previously published data cited in the text. MFL, CHD prepared the original draft. PB and CHD developed a software plugin introduced in this manuscript. MFL, WBS, JC, NPR, CHD contributed to the writing and editing of the manuscript. All authors approved the manuscript.

### Data Availability

Genotypic and phenotypic data will be made available at time of publication through the CIMMYT Dataverse repository using the standard data terms, with a permanent link and DOI, as is standard practice for CIMMYT. Examples can be viewed at data.cimmyt.org. Scripts will be made publicly available at time of publication as supplemental material on GitHub.

## References

Azodi CB, Bolger E, McCarren A, et al (2019) Benchmarking Parametric and Machine Learning Models for Genomic Prediction of Complex Traits. G3 (Bethesda) 9:3691–3702. 10.1534/g3.119.400498

Babu R, Rojas NP, Gao S, et al (2013) Validation of the effects of molecular marker polymorphisms in LcyE and CrtRB1 on provitamin A concentrations for 26 tropical maize populations. Theor Appl Genet 126:389–399. 10.1007/s00122-012-1987-3

Beck HE, Zimmermann NE, McVicar TR, et al (2018) Present and future Köppen-Geiger climate classification maps at 1-km resolution. Sci Data 5:180214. 10.1038/sdata.2018.214

Bernstein PS, Arunkumar R (2021) The emerging roles of the macular pigment carotenoids throughout the lifespan and in prenatal supplementation. Journal of Lipid Research 62:100038. 10.1194/jlr.TR120000956

Bhat JA, Ali S, Salgotra RK, et al (2016) Genomic Selection in the Era of Next Generation Sequencing for Complex Traits in Plant Breeding. Front Genet 7:. 10.3389/fgene.2016.00221

Blessin CW, Brecher JD, Dimler RJ, et al (1963) Carotenoids of Corn and Sorghum. III. Variation in Xanthophylls and Carotenes in Hybrid, Inbred, and Exotic Corn Lines. Cereal Chemistry

Bohn T, Desmarchelier C, El SN, et al (2019) β-Carotene in the human body: metabolic bioactivation pathways – from digestion to tissue distribution and excretion. Proceedings of the Nutrition Society 78:68–87. 10.1017/S0029665118002641

Bouis HE, Welch RM (2010) Biofortification-A Sustainable Agricultural Strategy for Reducing Micronutrient Malnutrition in the Global South. Crop Sci 50:S-20–S-32. 10.2135/cropsci2009.09.0531

Bradbury PJ, Zhang Z, Kroon DE, et al (2007) TASSEL: software for association mapping of complex traits in diverse samples. Bioinformatics 23:2633–2635. 10.1093/bioinformatics/btm308

Campos GDL, Gianola D, Rosa GJM, et al (2010) Semi-parametric genomic-enabled prediction of genetic values using reproducing kernel Hilbert spaces methods. Genetics Research 92:295–308. 10.1017/S0016672310000285

Clark SA, Van Der Werf J (2013) Genomic Best Linear Unbiased Prediction (gBLUP) for the Estimation of Genomic Breeding Values. In: Gondro C, Van Der Werf J, Hayes B (eds) Genome-Wide Association Studies and Genomic Prediction. Humana Press, Totowa, NJ, pp 321–330

Crossa J, Pérez P, Hickey J, et al (2014) Genomic prediction in CIMMYT maize and wheat breeding programs. Heredity 112:48–60. 10.1038/hdy.2013.16

Crossa J, Pérez-Rodríguez P, Cuevas J, et al (2017) Genomic Selection in Plant Breeding: Methods, Models, and Perspectives. Trends in Plant Science 22:961–975. 10.1016/j.tplants.2017.08.011

Diepenbrock CH, Ilut DC, Magallanes-Lundback M, et al (2021) Eleven biosynthetic genes explain the majority of natural variation in carotenoid levels in maize grain. The Plant Cell 33:882–900. 10.1093/plcell/koab032

Endelman JB (2011) Ridge Regression and Other Kernels for Genomic Selection with R Package rrBLUP. The Plant Genome 4:250–255. 10.3835/plantgenome2011.08.0024

Flint-Garcia SA, Thuillet A-C, Yu J, et al (2005) Maize association population: a high-resolution platform for quantitative trait locus dissection. Plant J 44:1054–1064. 10.1111/j.1365-313X.2005.02591.x

Friedman J, Hastie T, Tibshirani R (2010) Regularization Paths for Generalized Linear Models via Coordinate Descent. J Stat Soft 33:. 10.18637/jss.v033.i01

Galicia L, Nurit E, Rosales A, Palacios-Rojas N (2009) Maize nutrition quality and plant tissue analysis laboratory: laboratory protocols 2008. CIMMYT

Garnier, Simon, Ross, et al (2023) viridis(Lite) - Colorblind-Friendly Color Maps for R

Gianola D, Fernando RL, Stella A (2006) Genomic-Assisted Prediction of Genetic Value With Semiparametric Procedures. Genetics 173:1761–1776. 10.1534/genetics.105.049510

Guo R, Dhliwayo T, Mageto EK, et al (2020) Genomic Prediction of Kernel Zinc Concentration in Multiple Maize Populations Using Genotyping-by-Sequencing and Repeat Amplification Sequencing Markers. Front Plant Sci 11:534. 10.3389/fpls.2020.00534

Habier D, Fernando RL, Dekkers JCM (2007) The Impact of Genetic Relationship Information on Genome-Assisted Breeding Values. Genetics 177:2389–2397. 10.1534/genetics.107.081190

HarvestPlus, International Center for Tropical Agriculture (CIAT), Cali, Colombia, Andersson M (2017) Progress update: Crop development of biofortified staple food crops under HarvestPlus. AJFAND 17:11905–11935. 10.18697/ajfand.78.HarvestPlus05

Heslot N, Jannink J-L, Sorrells ME (2015) Perspectives for Genomic Selection Applications and Research in Plants. Crop Science 55:1–12. 10.2135/cropsci2014.03.0249

Heslot N, Yang H-P, Sorrells ME, Jannink J-L (2012) Genomic Selection in Plant Breeding: A Comparison of Models. Crop Science 52:146–160. 10.2135/cropsci2011.06.0297

Hodge C, Taylor C (2023) Vitamin A Deficiency. In: StatPearls. StatPearls Publishing, Treasure Island (FL)

Krinsky NI, Landrum JT, Bone RA (2003) BIOLOGIC MECHANISMS OF THE PROTECTIVE ROLE OF LUTEIN AND ZEAXANTHIN IN THE EYE. Annu Rev Nutr 23:171–201. 10.1146/annurev.nutr.23.011702.073307

Labroo MR, Rutkoski JE (2022) New cycle, same old mistakes? Overlapping vs. discrete generations in long-term recurrent selection. BMC genomics 23:. 10.1186/s12864-022-08929-3

Manjeru P, Van Biljon A, Labuschagne M (2019) The development and release of maize fortified with provitamin A carotenoids in developing countries. null 59:1284–1293. 10.1080/10408398.2017.1402751

Meuwissen TH, Hayes BJ, Goddard ME (2001) Prediction of total genetic value using genome-wide dense marker maps. Genetics 157:1819–1829

Monier B, Casstevens TM, Bradbury PJ, Buckler ES (2022) rTASSEL: An R interface to TASSEL for analyzing genomic diversity. Journal of Open Source Software 7:4530. 10.21105/joss.04530

Montesinos López OA, Montesinos López A, Crossa J (2022) Reproducing Kernel Hilbert Spaces Regression and Classification Methods. In: Montesinos López OA, Montesinos López A, Crossa J (eds) Multivariate Statistical Machine Learning Methods for Genomic Prediction. Springer International Publishing, Cham, pp 251–336

Ogutu JO, Schulz-Streeck T, Piepho H-P (2012) Genomic selection using regularized linear regression models: ridge regression, lasso, elastic net and their extensions. BMC Proceedings 6:S10. 10.1186/1753-6561-6-S2-S10

Ortiz D, Ponrajan A, Bonnet JP, et al (2018) Carotenoid Stability during Dry Milling, Storage, and Extrusion Processing of Biofortified Maize Genotypes. J Agric Food Chem 66:4683– 4691. 10.1021/acs.jafc.7b05706

Owens BF, Lipka AE, Magallanes-Lundback M, et al (2014) A Foundation for Provitamin A Biofortification of Maize: Genome-Wide Association and Genomic Prediction Models of Carotenoid Levels. Genetics 198:1699–1716. 10.1534/genetics.114.169979

Ozimati A, Kawuki R, Esuma W, et al (2019) Genetic Variation and Trait Correlations in an East African Cassava Breeding Population for Genomic Selection. Crop Science 59:460–473. 10.2135/cropsci2018.01.0060

Palacios-Rojas N, Twumasi-Afriye S, Friesen DK, et al (2017) Lineamientos para el control de calidad de semilla y grano de maíz de alta calidad proteica (QPM): experiencia en el desarrollo y promoción de QPM en Latinoamérica. CIMMYT

Paula RG de, Gabriela S. Pereira, Igor G. de Paula, et al (2020) Multipopulation recurrent selection: An approach with generation and population effects in selection of self- pollinated progenies. Agronomy journal 112:4602–4612. 10.1002/agj2.20422

Perez P, Campos G de los (2014) Genome-Wide Regression and Prediction with the BGLR Statistical Package. Genetics 198:483–495

Pixley K, Rojas NP, Babu R, et al (2013) Biofortification of Maize with Provitamin A Carotenoids. In: Tanumihardjo SA (ed) Carotenoids and Human Health. Humana Press, Totowa, NJ, pp 271–292

Prasanna BM, Palacios-Rojas N, Hossain F, et al (2020) Molecular Breeding for Nutritionally Enriched Maize: Status and Prospects. Frontiers in Genetics 10:

R Core Team (2021) R: A Language and Environment for Statistical Computing. R Foundation for Statistical Computing, Vienna, Austria

Rakotondramanana M, Tanaka R, Pariasca-Tanaka J, et al (2022) Genomic prediction of zinc-biofortification potential in rice gene bank accessions. Theor Appl Genet 135:2265–2278. 10.1007/s00122-022-04110-2

Saltzman A, Birol E, Bouis HE, et al (2013) Biofortification: Progress toward a more nourishing future. Global Food Security 2:9–17. 10.1016/j.gfs.2012.12.003

Suwarno WB, Pixley KV, Palacios-Rojas N, et al (2015) Genome-wide association analysis reveals new targets for carotenoid biofortification in maize. Theor Appl Genet 128:851– 864. 10.1007/s00122-015-2475-3

Tibbs-Cortes LE, Guo T, Li X, et al (2022) Genomic prediction of tocochromanols in exotic-derived maize. The Plant Genome e20286. 10.1002/tpg2.20286

Tibshirani R (1996) Regression Shrinkage and Selection via the Lasso. Journal of the Royal Statistical Society Series B (Methodological) 58:267–288

Usai MG, Goddard ME, Hayes BJ (2009) LASSO with cross-validation for genomic selection. Genet Res (Camb) 91:427–436. 10.1017/S0016672309990334

Wang J, Zhang Z (2021) GAPIT Version 3: Boosting Power and Accuracy for Genomic Association and Prediction. Genomics, Proteomics & Bioinformatics 19:629–640. 10.1016/j.gpb.2021.08.005

Weber EJ (1987) Carotenoids and tocols of corn grain determined by HPLC. J Am Oil Chem Soc 64:1129–1134. 10.1007/BF02612988

Wickham H (2016) ggplot2: Elegant Graphics for Data Analysis. Springer-Verlag New York

Windhausen VS, Atlin GN, Hickey JM, et al (2012) Effectiveness of Genomic Prediction of Maize Hybrid Performance in Different Breeding Populations and Environments. G3 (Bethesda) 2:1427–1436. 10.1534/g3.112.003699

Wirth JP, Petry N, Tanumihardjo SA, et al (2017) Vitamin A Supplementation Programs and Country-Level Evidence of Vitamin A Deficiency. Nutrients 9:190. 10.3390/nu9030190

Woodhouse MR, Cannon EK, Portwood JL, et al (2021) A pan-genomic approach to genome databases using maize as a model system. BMC Plant Biol 21:385. 10.1186/s12870-021-03173-5

World Health Organization (2009) Global prevalence of vitamin A deficiency in populations at risk 1995-2005: WHO global database on vitamin A deficiency. 55

World Health Organization (2014) Xerophthalmia and night blindness for the assessment of clinical vitamin A deficiency in individuals and populations

Yan J, Kandianis CB, Harjes CE, et al (2010) Rare genetic variation at Zea mays crtRB1 increases β-carotene in maize grain. Nat Genet 42:322–327. 10.1038/ng.551

Zou H, Hastie T (2005) Regularization and Variable Selection via the Elastic Net. Journal of the Royal Statistical Society Series B (Statistical Methodology) 67:301–320

